# Structure of a bacterial ATP synthase

**DOI:** 10.1101/463968

**Authors:** Hui Guo, Toshiharu Suzuki, John L. Rubinstein

## Abstract

ATP synthases produce ATP from ADP and inorganic phosphate with energy from a transmembrane proton motive force. Bacterial ATP synthases have been studied extensively because they are the simplest form of the enzyme and because of the relative ease of genetic manipulation of these complexes. We expressed the *Bacillus* PS3 ATP synthase in *Eschericia coli*, purified it, and imaged it by cryo-EM, allowing us to build atomic models of the complex in three rotational states. The position of subunit *ε* shows how it is able to inhibit ATP hydrolysis while allowing ATP synthesis. The architecture of the membrane region shows how the simple bacterial ATP synthase is able to perform the same core functions as the equivalent, but more complicated, mitochondrial complex. The structures reveal the path of transmembrane proton translocation and provide a model for understanding decades of biochemical analysis interrogating the roles of specific residues in the enzyme.

## Introduction

Adenosine triphosphate (ATP) synthases are multi-subunit protein complexes that use an electrochemical proton motive force across a membrane to make the cell’s supply of ATP from adenosine diphosphate (ADP) and inorganic phosphate (Pi). These enzymes are found in bacteria and chloroplasts as monomers, and in mitochondria as rows of dimers that bend the inner membrane to facilitate formation of the mitochondrial cristae ^1,2^. Proton translocation across the membrane-embedded F_O_ region of the complex occurs via two offset half-channels ^3,4^. Studies with *Bacillus* PS3 ATP synthase in liposomes showed that proton translocation may be driven by ΔpH or ΔΨ alone ^5^. The passage of protons causes rotation of a rotor subcomplex, inducing conformational change in the catalytic F_1_ region to produce ATP ^6^ while a peripheral stalk subcomplex holds the F_1_ region stationary relative to the spinning rotor during catalysis. For the mitochondrial enzyme, X-ray crystallography has been used to determine structures of the soluble F_1_ region ^7^, partial structures of the peripheral stalk subcomplex alone ^8^ and with the F_1_ region ^9^, and structures of the F_1_ region with the membrane-embedded ring of *c*-subunits attached ^10,11^. Recent breakthroughs in electron cryomicroscopy (cryo-EM) allowed the structures of the membrane-embedded F_O_ regions from mitochondrial and chloroplast ATP synthases to be determined to near-atomic resolutions ^12–15^.

Compared to their mitochondrial counterparts, bacterial ATP synthases have a simpler subunit composition. The F_1_ region consists of subunits *α_3_β_3_γδε*, while the F_O_ region is usually formed by three subunits with the stoichiometry *ab_2_c_9-15_*. Chloroplasts and a few bacteria, such as *Paracoccus denitrificans*, possess two different but homologous copies of subunit *b*, named subunits *b* and *b′* ^6^. Each copy of subunit *α* and *β* contains a nucleotide binding site. The non-catalytic *α* subunits each bind to a magnesium ion (Mg^2+^) and a nucleotide, while the catalytic *β* subunits can adopt different conformations and bind to Mg-ADP (*β_DP_*), Mg-ATP (*β_TP_*), or remain empty (*β_E_*). Crystal structures of bacterial F_1_-ATPases and *c*-rings from the F_O_ regions of several species have been determined ^16–24^. Structures of intact ATP synthases from *E. coli* have been determined to overall resolutions of 6 to 7 Å by cryo-EM, with the F_O_ region showing lower quality than the rest of the maps, presumably due to conformational flexibility ^25^. In structures of both intact ATP synthase ^25^ and dissociated F_1_-ATPase ^17,19^ from bacteria, subunit *ε* adopts an “up” conformation that inhibits the ATP hydrolysis by the enzyme. In the thermophilic bacterium *Bacillus* PS3, this subunit *ε* mediated inhibition is dependent on the concentration of free ATP ^26–28^. Low ATP concentrations (*eg*. < 0.7 mM) promote the inhibitory *up* conformation while a permissive “down” conformation can be induced by a high concentration of ATP (*eg*. > 1 mM). This mechanism would allow the *Bacillus* PS3 ATP synthase to run in reverse, establishing a proton motive force by ATP hydrolysis, when the ATP concentration is sufficient to do so without depleting the cell’s supply of ATP. In *E. coli*, however, in the absence of a sufficient proton motive force to drive ATP synthesis, inhibition of ATP hydrolysis by subunit *ε* persists even when the concentration of free ATP is high ^29,30^.

Although bacterial ATP synthases have been subjected to extensive biochemical analysis, high-resolution structural information is lacking for the intact enzyme or the membrane-embedded proton-conducting subunit *a* and the associated subunit *b*. We determined structures of intact ATP synthase from *Bacillus* PS3 in three rotational states by cryo-EM. The structures reached overall resolutions of 2.9 to 3.1 Å (Fig. 1), allowing construction of nearly complete atomic models for the entire complex. The structures reveal how loops in subunit *a* of the bacterial enzyme fill the role of additional subunits in the F_O_ region of the mitochondrial enzyme. Most significantly, the structures provide a framework for understanding decades of mutagenesis experiments designed to probe the mechanism of ATP synthases.

**Figure 1.**
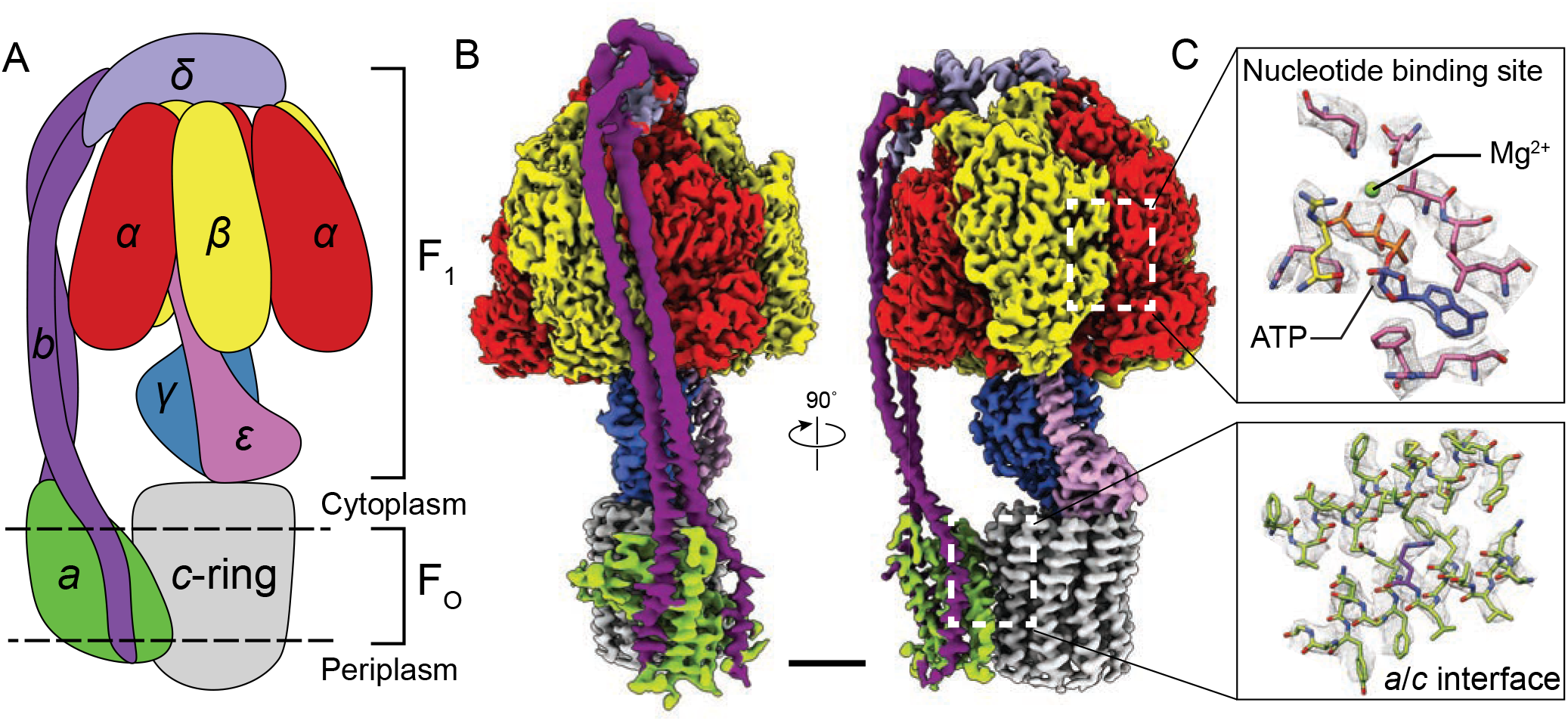
Overall structure of *Bacillus* PS3 ATP synthase. **A,** Cartoon of ATP synthase. **B,** Cryo-EM map of ATP synthase with subunits coloured the same as the cartoon. **C,** Example map density that allowed construction of an atomic model. Scale bar, 30 Å.

## Results and discussions

### Structure determination and overall architecture

Subunits of *Bacillus* PS3 ATP synthase, including subunit *β* bearing an N-terminal 10×His tag, were expressed from a plasmid in *E. coli* strain DK8, which lacks endogenous ATP synthase ^31,32^. The complex was extracted from membranes with detergent, purified by metal-affinity chromatography, and subjected to cryo-EM analysis (Fig. 1 - figure supplement 1). Three conformations corresponding to different rotational states of the enzyme were identified by abinitio 3D classification and refined to high resolution. The 3D classes contain 45, 35, and 20 % of particle images and the overall resolutions of the corresponding cryo-EM maps were 2.9, 2.9, and 3.1 Å, respectively (Fig. 1 - figure supplements 2 and 3). Estimation of local resolution suggests that the F_1_ regions of the maps, which are larger than the F_O_ regions and appear to dominate the image alignment process, are mostly at between 2.5 and 3.5 Å resolution, whereas the F_O_ regions were limited to lower resolution (Fig. 1 - figure supplement 3). Focused refinement ^33^ of the F_O_ region and peripheral stalk subunits *ab_2_c_10_* and *δ* (corresponding to the subunit *OSCP* in mitochondrial ATP synthase) improved the resolution of the F_O_ regions considerably for all three classes but not enough to resolve density for most of the amino acid side chains. An improved map of the F_O_ region was obtained by focused refinement of the membrane-embedded region only, excluding the soluble portion of subunit *b* with particle images from all three classes (Fig. 1 - figure supplement 2). Overall, amino acid side chain detail can be seen for subunits *α_3_*, *α_3_*, *γ*, *δ*, *ε*, *a*, *c_10_*-ring, and the transmembrane *α* -helices of *b_2_* (Fig.1 - figure supplements 4). The soluble region of the two *b*-subunits was modeled as poly-alanine.

The general architecture of the enzyme resembles *E. coli* ATP synthase ^25^ and the more distantly related spinach chloroplast enzyme ^15^ but with striking differences. As observed previously in a *Bacillus* PS3 F_1_-ATPase crystal structure (PDB 4XD7) ^19^, the three catalytic *β* subunits adopt “open”, “closed”, and “open” conformations, different from the “half-closed”, “closed”, and “open” conformations seen in the auto-inhibited *E. coli* F_1_-ATPase ^17^, and the “closed”, “closed”, and “open” conformations seen in chloroplast ATP synthase ^15^ and most mitochondrial ATP synthase structures ^7,10^. This difference, with the *half-closed β_DP_* of the *E. coli* enzyme appearing as *open* in the *Bacillus* PS3 enzyme, suggests species-specific differences in inhibition by subunit *ε* (Fig. 1B, pink density), which inserts into the *α*/*β* interface and forces *β_DP_* into the *open* conformation.

In the F_O_ region, one copy of subunit *b* is positioned at a location equivalent to that of the mitochondrial subunit *b*, while the second copy occupies the position of yeast subunit *8* (mammalian *A6L*) on the other side of subunit *a* (Fig. 1B). Despite the different *c*-ring sizes (10 *c*-subunits in *Bacillus* PS3 versus 14 in spinach chloroplasts), the backbone positions of subunits *ab_2_* from *Bacillus* PS3 overlap with subunits *abb′* from spinach chloroplast ATP synthase ^15^ (Fig. 1 - figure supplement 5A). Comparison of the atomic model of the F_O_ region from *Bacillus* PS3 and the backbone model of the *E. coli* complex from cryo-EM at ~7 Å resolution (PDB 5T4O) ^25^ showed significant structural differences in transmembrane α-helices of subunit *b* relative to subunit *a* (Fig. 1 - figure supplement 5B). Rather than reflecting true differences between *E. coli* and *Bacillus* PS3 ATP synthase structures, these deviations likely suggest that the 6 to 8 Å resolution *E. coli* maps were not at sufficient resolution to allow accurate backbone tracing of F_O_ subunits.

### Flexibility in the peripheral and central stalks

As expected, the most striking difference between the three rotational states of the *Bacillus* PS3 structure is the angular position of the rotor (subunits *γεc_10_*) (Fig. 2A, Video 1). The structure of the ATP synthase, with three *αβ* pairs in the F_1_ region and ten *c*-subunits in the F_O_ region, results in symmetry mismatch between the 120° steps of the F_1_ motor and 36° steps of the F_O_ motor. The 120° steps of the F_1_ motor gives an average rotational step of 3.3 *c*-subunits, with the closest integer steps being 3, 4 and 3 *c*-subunits. By comparing the positions of equivalent *c*-subunits in different rotational states, the observed rotational step sizes in the three rotational states of the ATP synthase appear to be almost exactly 3, 4 and 3 *c*-subunits (Fig. 2B). At the present resolution, the structures of subunit *a* and the *c-*ring do not appear to differ between rotary states. Similar integer step sizes were found in yeast ATP synthase ^34^ and V-ATPase ^35^, which also contain 10 *c*-subunits. However, non-integer steps were seen in the chloroplast (14 *c*-subunits) ^15^ and bovine (8 *c*-subunits) ^36^ ATP synthases, indicating that the *c*-subunit steps between the rotational states of rotary ATPases likely depends on the number of *c*-subunits.

**Figure 2.**
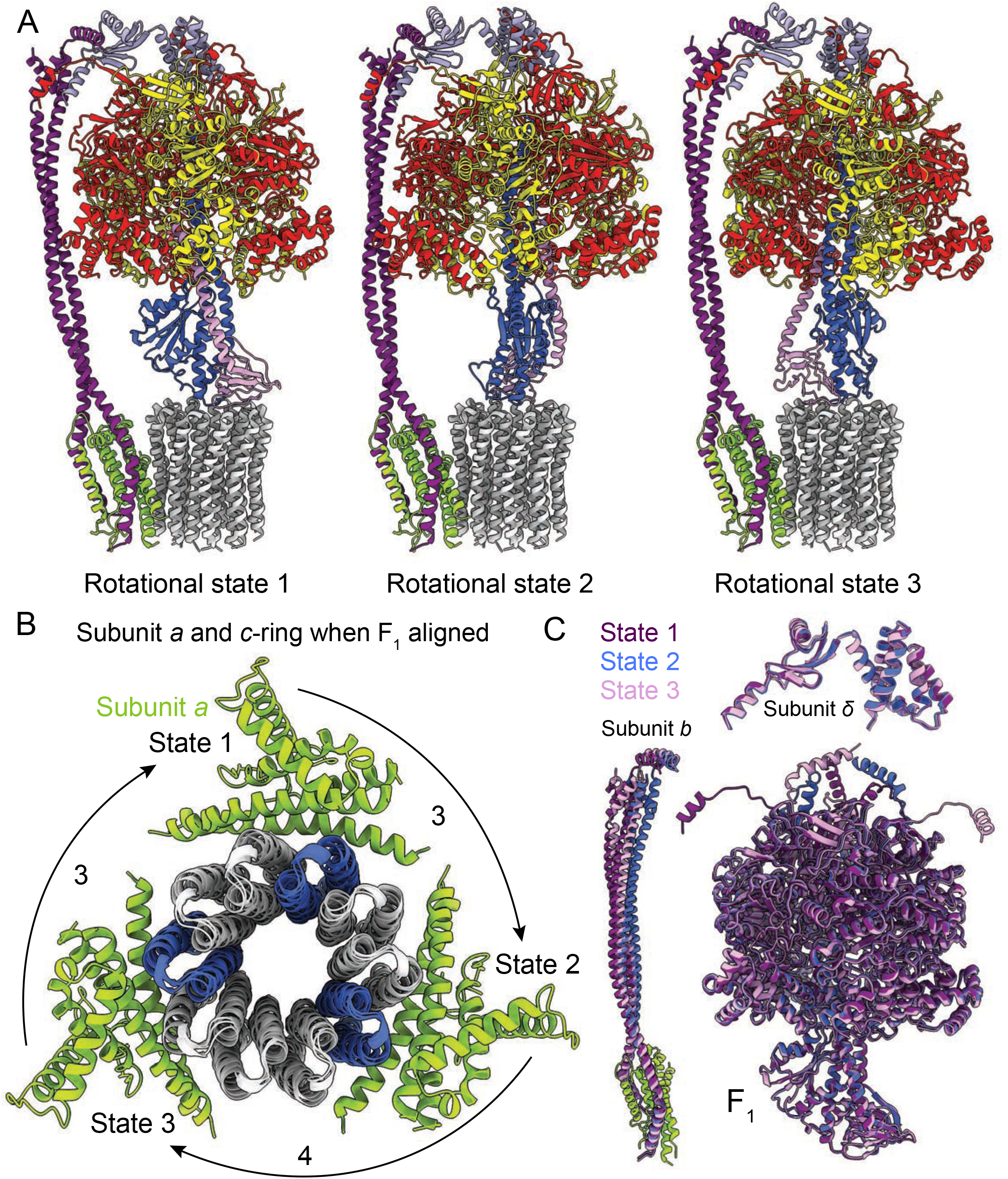
Rotational states of ATP synthase. **A,** Atomic models of the three rotational states of *Bacillus* PS3 ATP synthase with subunits coloured the same as in Figure 1. **B,** Top view of the *c*-ring and subunit *a* of the three rotational states from the cytoplasm when the F_1_ regions of the three states are aligned. Rotation steps of the complex between states are ~3, 4, and 3 *c*-subunits. **C,** Comparison of the atomic models of subunits *b*, *δ*, and other F_1_ region subunits in the different rotational states. The *b* subunits appear to be the most flexible part of the enzyme.

Flexibility is thought to be important for the smooth transmission of power between the F_1_ and F_O_ regions, which often have mismatched symmetries ^37–39^. Earlier studies suggested that the central stalk (subunits *γ* and *ε* in bacteria) is the main region responsible for the transient storage of torsional energy in rotary ATPases ^40,41^. Comparison of the three rotational states of the *Bacillus* PS3 enzyme also shows that C-terminal water-soluble part of subunit *b* displays the most significant conformational variability between states, while the subunits in the F_1_ region show little flexibility beyond the catalytic states of the *αβ* pairs (Fig. 2C; Video 1). The structure of the yeast ATP synthase F_O_ dimer ^12^, which lacked the the F_1_ region and an intact peripheral stalk, showed that the *c*-ring and subunit *a* are held together by hydrophobic interactions rather than by the peripheral stalk. In *Bacillus* PS3 ATP synthase, the peripheral stalk is structurally simpler and more flexible than in yeast mitochondria ^14^, suggesting that the bacterial subunits *a* and the *c*-ring are also held together by hydrophobic interactions and not the peripheral stalk. Given that these structures represent resting states of the bacterial ATP synthase, additional subunits, such as those in the central stalk, may show flexibility while under strain during rotation.

### Nucleotide binding in the F_1_ region and inhibition by subunit ε

The structure of the F_1_ region of the intact *Bacillus* PS3 ATP synthase and the earlier crystal structure of the dissociated F_1_-ATPase (PDB 4XD7) ^19^ both show that the three catalytic *β*-subunits (*β_E_*, *β_TP_*, and *β_D_*_P_) adopt “open”, “closed”, and “open” conformations, respectively (Fig. 3A). In the crystal structure, which was prepared in the presence of CyDTA (trans-1,2-Diaminocyclohexane-N, N, N*′*, N*′*-tetraacetic acid monohydrate) as a chelating agent, there was no nucleotide in the three noncatalytic sites of the three *α-*subunits and the only nucleotide in a catalytic site was an ADP molecule without a Mg^2+^ ion in the *β_TP_* site. In contrast, all three non-catalytic sites in the cryo-EM map are occupied by Mg-ATP, while a Mg-ADP molecule and a weak density tentatively assigned to Pi are found in the *β_TP_* site and by the p-loop of *β_E_*, respectively. The presence of physiological Mg^2+^ ions and nucleotide occupancy ^42^ in the cryo-EM map suggest that it shows a snapshot of the enzyme in the middle of its physiological catalytic cycle.

**Figure 3.**
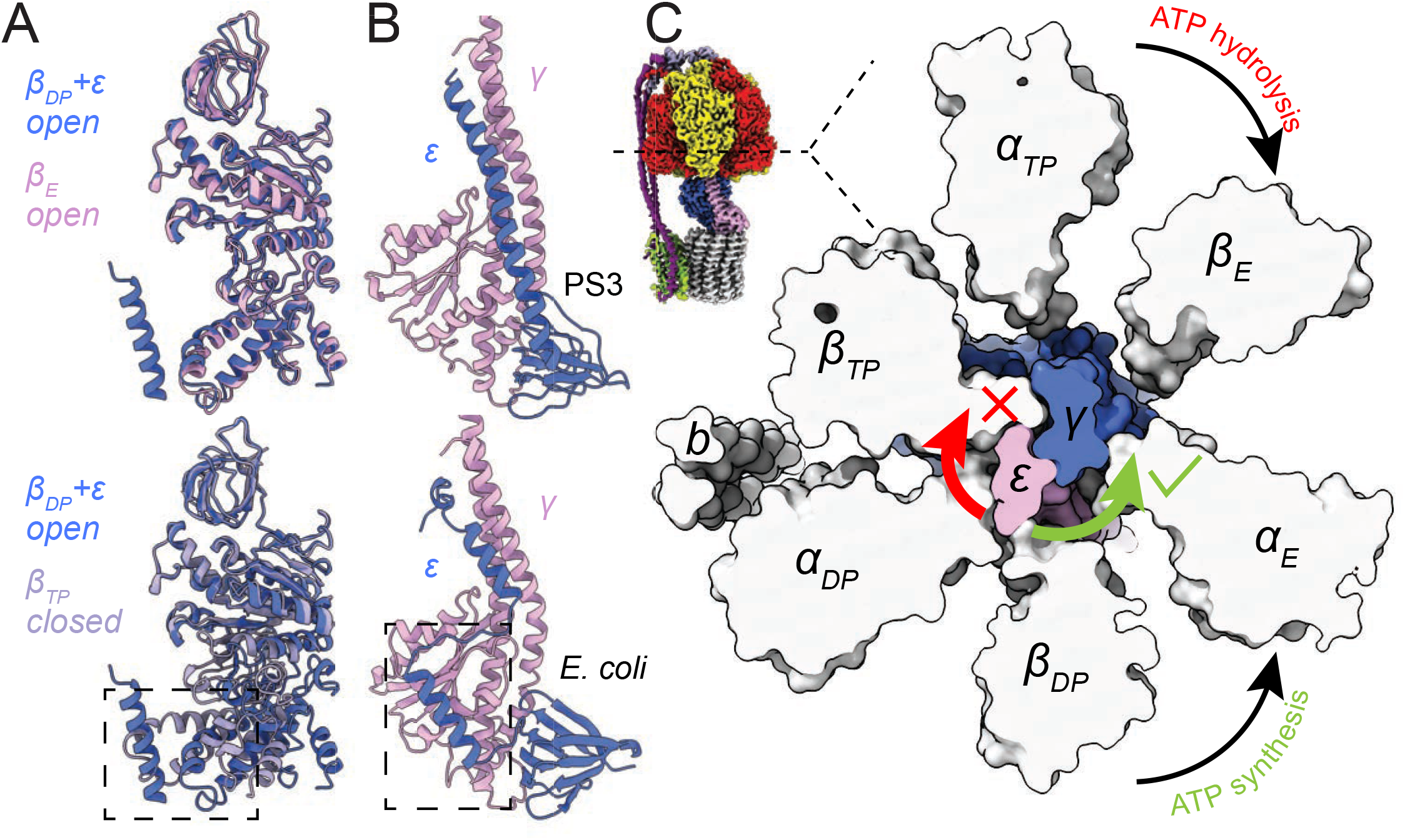
Inhibition of ATP hydrolysis by subunit *ε*. **A,** Comparison of *β_DP_* (blue) with *β_E_* (pink, top) and *β_TP_* (light purple, bottom). *β_DP_* is forced to adopt an *open* conformation by subunit *ε* (dashed box). **B,** Comparison of subunits *γ* (pink) and *ε* (blue) of ATP synthases from *Bacillus* PS3 (top) and *E. coli* (bottom, PDB 3OAA ^17^). The dashed box shows additional interaction between subunits *ε* and *γ* in the *E. coli* complex. **C,** Cross section through the catalytic F_1_ region of the *Bacillus* PS3 ATP synthase. Subunit *ε* (pink) in the rotor is blocked from rotating in the direction of ATP hydrolysis (clockwise) by *β_TP_* but is free to rotate in the direction of ATP synthesis (counterclockwise).

*Bacillus* PS3 ATP synthase is found in a conformation where ATP synthesis is permitted but ATP hydrolysis is auto-inhibited. In this state subunit *ε* maintains an *up* conformation and inserts into the *α_DP_β_DP_* interface, forcing *β_DP_* to adopt an open conformation (Fig. 3A, lower, dashed box) ^19^. In the crystal structure (PDB 4XD7) ^19^, the C-terminal sequence of subunit *ε* was modeled as two α-helical segments broken at Ser 106, while the cryo-EM structures show the C-terminal part is in fact entirely α-helical. In comparison, subunit *ε* from the auto-inhibited *E. coli* F_1_-ATPase structure (PDB 3OAA) ^17^ maintains its two C-terminal α-helices (Fig. 3B), with its *β_DP_* adopting a *half-closed* conformation that binds to Mg-ADP. The C-terminal α-helix of the *E. coli* subunit α inserts slightly deeper into the *α_DP_β_DP_* interface but overall in a manner similar to that of the *Bacillus* PS3 subunit *ε*. However, the second *α*-helix in *E. coli* is offset by a ten-residue loop that allows it to interact with subunit *γ*. This interaction (Fig. 3B, lower, dashed box) may stabilize the *up* conformation of subunit *ε* in *E. coli*, explaining why auto-inhibition in *E. coli* does not depend on ATP concentration ^29,30^ while in *Bacillus* PS3 it does. Interestingly, during ATP synthesis, *Bacillus* PS3 subunit *ε* maintains the *up* conformation ^27^, suggesting that it only blocks ATP hydrolysis but not ATP synthesis. For a canonical ATP synthase, the substrates ADP and Pi bind to an *open β_E_*. The *β_E_* subsequently transitions to become the *closed β_DP_* and then *β_TP_*, driven by rotation of the central rotor, producing an ATP molecule that is ultimately released when the *closed β_TP_* converts back to an open *β_E_* ^7^. For the *Bacillus* PS3 ATP synthase to produce ATP with subunit *ε* in the *up* conformation, substrate would need to bind to the *β_DP_* site instead of the usual *β_E_* site, with an ATP molecule produced on transition to a *closed β_T_*_P_. The cryo-EM maps show that a clash between subunit *ε* and *β_T_*_P_ blocks the central rotor turning in the direction of ATP hydrolysis while it is still free to turn in the direction of ATP synthesis (Fig. 3C), explaining the ability of subunit *ε* to selectively inhibit ATP hydrolysis ^27^.

### Subunit organization in the F_O_ region

In the bacterial ATP synthase structure, the F_O_ subunits *ab_2_* display an organization similar to the yeast F_O_ complex (PDB 6B2Z, Fig. 4A) ^12^. Subunit *a* and the first copy of subunit *b* occupy the same positions as their yeast counterparts, while the second copy of subunit *b* is found at a position equivalent to subunit *8* in the yeast enzyme, which is known as *A6L* in mammals. Atomic models for ATP synthase from mitochondria ^12–14^ and chloroplasts ^15^ support the idea that transmembrane proton translocation in ATP synthases occurs via two offset half-channels formed by subunit *a* ^3,4^. Subunit *a* from *Bacillus* PS3 shares 21.0% and 29.1% sequence identity with its yeast and chloroplast homologs, respectively, and the atomic model shows that the folding of these homologs is mostly conserved (Fig. 4 - figure supplement 1). Multi-sequence alignment of subunit *a* from different species indicates that bacterial and chloroplast subunit *a* contain a larger periplasmic loop between *α*-helices 3 and 4 than found in the mitochondrial subunit (Fig. 4A, left; Fig. 4 - figure supplement 2). The sequence for this loop varies significantly among species, suggesting that it is unlikely to be involved in the core function of proton translocation, despite being proximal to the cytosolic proton half-channel. Yeast and mammalian mitochondrial ATP synthases contain subunit *f*, which has a transmembrane α-helix adjacent to the transmembrane α-helix 1 of subunit *a* (Fig. 4A, right), anchoring subunit *b* between α-helices 5 and 6 of subunit *a*. The location of the loop between α-helices 3 and 4 of the *Bacillus* PS3 subunit *a* suggests that it serves a similar structural role, compensating for the lack of subunit *f* in bacteria. The loop forms an additional interface with subunit *b* near the periplasmic side of the membrane region and may interact with the N terminus of subunit *b* in the periplasm as well. Two interfaces are also present between the second copy of subunit *b* and subunit *a*, one with the first transmembrane α-helix, and the other with the hairpin of α-helices 3 and 4 (Fig. 4A). The structure suggests that two interfaces are necessary for subunits *a* and b to maintain a stable interaction.

**Figure 4.**
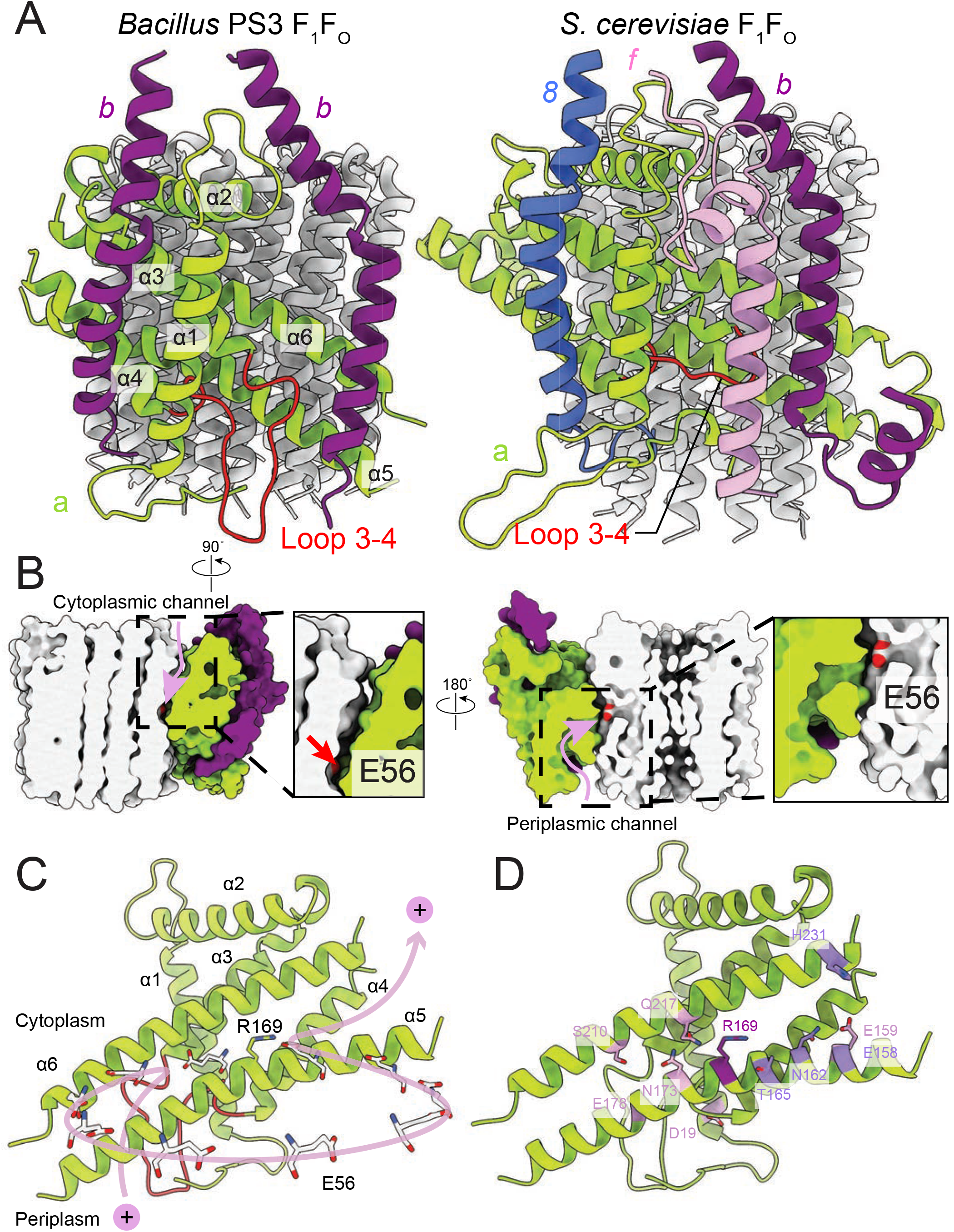
F_O_ region subunits and proton translocation in *Bacillus* PS3 ATP synthase. **A,** Comparison of the F_O_ regions from *Bacillus* PS3 (left) and *S. cerevisiae* (right). **B**, Cross sections through a surface representation of the F_O_ region (simulated with rolling of a 1.4 Å sphere ^43^) show the cytoplasmic (left) and periplasmic (right) proton half-channels. **C,** Proton translocation pathway of *Bacillus* PS3 ATP synthase. **D,** Subunit *a* of *Bacillus* PS3 ATP synthase. Arg 169 is in purple, important residues for proton translocation identified by mutagenesis in *E. coli* ATP synthase are in pink, and other residues that appears to contribute to proton transfer in the cytosolic proton half-channel are in light purple.

### Proton translocation through the F_O_ region

The *Bacillus* PS3 ATP synthase structure implies a path for proton translocation through the bacterial complex involving two half-channels and similar to the paths described for the mitochondrial and chloroplast enzymes. The cytoplasmic half-channel consists of an aqueous cavity at the interface of subunit *a* and the *c*-ring (Fig. 4B, left). The periplasmic half-channel is formed from a cavity between α-helices 1, 3, 4 and 5 of subunit *a*, and reaches the *c*-ring via a gap between α-helices 5 and 6 (Fig. 4B, right). In the atomic model, both channels are visible when modelling the surface with a 1.4 Å sphere that mimics a water molecule ^43^ (Fig. 4B). The channels are wide and hydrophilic, suggesting that water molecules could pass freely through each of the channels before accessing the conserved Glu 56 of the *c*-subunits. During ATP synthesis, protons travel to the middle of the *c*-ring via the periplasmic half-channel and bind to the Glu 56 residue of a subunit *c* (Fig. 4C). Protonation of the glutamate allows rotation of the ring counter-clockwise, when viewed from F_1_ towards F_O_, delivering the subunit *c* into the hydrophobic lipid bilayer. Protonation of the remaining nine subunits in the *c*-ring returns the first glutamate to subunit *a*, now into the cytoplasmic half-channel, where it releases its proton to the cytoplasm due to interaction with the positively charged Arg 169 of subunit *a*.

In eukaryotes, subunit *a* is encoded by the mitochondrial genome, limiting genetic interrogation of the roles of different residues. In contrast, numerous mutagenesis studies have been performed on bacterial subunits *a* and *b*, with *E. coli* ATP synthase being the most frequently studied ^44,45^. A single G9D mutation in the *E. coli* subunit *b* (positionally equivalent to Y13D in *Bacillus* PS3), results in assembled but non-functional ATP synthase ^46^, while multiple N-terminal mutations in subunit *b* can either disrupt enzyme assembly or ATP hydrolysis ^47^. In *Bacillus* PS3, Tyr 13 is part of the transmembrane α-helix of subunit *b* and is adjacent to Gly 188 of subunit *a* (Fig. 4 - figure supplement 3, dashed box). In *E. coli* subunit a, Gly 188 is replaced by a leucine (Leu 229). Therefore, the G9D mutation in *E. coli* not only introduces a charged residue into a hydrophobic transmembrane α-helix, but also creates a steric clash with Leu 229 of subunit *a*, explaining why the mutation leads to an inactive enzyme. Remarkably, the single N-terminal membrane-embedded α-helix in each of the two copies of subunit *b* in the *Bacillus* PS3 ATP synthase forms different interactions with subunit *a* (Fig. 4A). One surface interacts with transmembrane α-helices 1, 2, 3, and 4 of subunit *a* while the other interacts with α-helices 5 and 6 and the loop between α-helices 3 and 4 of subunit *a*. Given that the N-terminal α-helix of subunit *b* makes interactions with different regions of subunit *a*, it is not surprising that mutations in this region are often detrimental to the assembly and activity of the complex. Cross-linking experiments suggested that the N terminus of the two copies of subunit *b* are in close proximity with each other ^48^. However, the atomic model shows that the transmembrane α-helix of the *b*-subunits are on opposite sides of subunit *a*, suggesting that the cross-linking results may be due to non-specific interactions of *b*-subunits from neighboring ATP synthases.

In *E. coli*, Arg 210 of subunit *a* (Arg 169 in *Bacillus* PS3) tolerates the fewest mutations ^49–52^. Recent structures of rotary ATPases suggest that the importance of this residue derives from its role in releasing protons bound to the Glu residues of the *c*-subunits as they enter the cytoplasmic half-channel, as well as preventing short-circuiting of the proton path by protons flowing between half-channels without rotation of the *c*-ring ^18,35,36,53,54^. Other residues in the *E. coli* subunit *a* identified by mutation as being functionally important include Glu 196 (Glu 159 in *Bacillus* PS3) ^55,56^, Glu 219 (Glu 178) ^55–57^, His 245 (Ser 210) ^51,58,59^, Asp 44 (Asp 19) ^60^, Asn 214 (Asn 173) ^49^, and Gln 252 (Gln 217) ^57,61^ (Fig. 4D). When mapped to the *Bacillus* PS3 structure, only Glu 196 (Glu 159 in *Bacillus* PS3) is close to the cytoplasmic half-channel. Extensive mutations of *E. coli* Glu 196 showed that enzyme activity depends on the charge and polarity of the residue with Glu > Asp > Gln = Ser = His > Asn > Ala > Lys ^55^. Therefore, the negative surface charge from Glu 196 (Glu 159) near the cytoplasmic half-channel facilitates proton transport across the lipid bilayer. The atomic model of subunit *a* also suggests that other residues such as *Bacillus* PS3 Thr 165, Asn 162, Glu 158, Tyr 228, and His 231, which are close to the cytoplasmic half-channel, may contribute to channel formation. Many functional residues identified by mutagenesis are clustered around the periplasmic half-channel. In the atomic model of the *Bacillus* PS3 subunit *a*, Asp 19 and Glu 178 are close to the periplasm, while Ser 210, Asn 173, and Gln 217 are deeper inside the membrane. Among these residues, Glu 178 and Ser 210 are considered to be more important to enzyme function than Asn 173 and Gln 217, as mutations of corresponding residues in *E. coli* are more likely to abolish the proton translocation by the complex ^44^. Interestingly, although many of these functional residues appear important, their mutation to amino acids that cannot be protonated or deprotonated often does not completely abolish proton translocation ^44^. The atomic model of *Bacillus* PS3 subunit *a* shows that the proton half-channels are wide enough for water molecule to pass through freely. This observation suggests that the function of these conserved polar and charged residues is not the direct transfer of protons during translocation. Rather, their presence may help maintain a hydrophilic environment for water-filled proton channels. This role allows different species to use unique sets of polar and charged residues forming their proton half-channels. For instance, the function of the Glu 219/His 245 pair in *E. coli* ^59^ is replaced by a Glu 178/Ser 210 pair in *Bacillus* PS3, and a His 185/Glu223 pair in yeast and human ATP synthases (Fig. 4 - figure supplement 2). This variability suggests a remarkably flexible proton translocation mechanism for this highly efficient macromolecular machine.

## Acknowledgements

We thank Dr. Samir Benlekbir (the Hospital for Sick Children) for helping with cryo-EM data collection and Prof. Tomitake Tsukihara (Osaka University, Japan) for discussions. This work was supported by Canadian Institutes of Health Research operating grant MOP 81294. Cryo-EM data was collected at the Toronto High-Resolution High-Throughput cryo-EM facility, supported by the Canada Foundation for Innovation and Ontario Research Fund. HG was supported by an Ontario Graduate Scholarship and a University of Toronto Excellence Award. TS was supported by Japan Society for the Promotion of Science Grants-in-Aid for Scientific Research (KAKENHI) Grant JP18H02409. JLR was supported by the Canada Research Chairs program. Cryo-EM maps are deposited in the Electron Microscopy Data Bank (EMD-XXXX and EMD-XXXX) and Protein Data Bank (PDB-XXX and PDB-XXX).

## Supplementary materials: Table 1

**Figure 1 - figure supplement 1-4**

**Figure 4 - figure supplement 1-3**

**Video 1**

### Material and Methods

#### Protein expression and purification

*E. coli* strain DK8, in which the genes encoding endogenous ATP synthase subunits were deleted ^31^, was transformed with plasmid pTR-ASDS ^32^ encoding *Bacillus* PS3 ATP synthase with a 10× His tag at the N terminus of subunit *β*. Transformed E. coli cells were grown in 2×TY medium at 37 °C for 20 hours before being harvested by centrifugation at 5,400 g. Cell pellets were resuspended in lysis buffer (50 mM Tris-HCl pH 7.4, 5 mM MgCl_2_, 10 % [w/v] glycerol, 5 mM 6-aminocaproic acid, 5 mM benzamidine, 1 mM PMSF) and lysed with three passes through an EmulsiFlex-C3 homogenizer (Avestin) at 15 to 20 kbar. All protein preparation steps were performed at 4 °C unless otherwise stated. Cell debris was removed at 12,250 g for 20 min and the cell membrane fraction was collected by centrifugation at 184,000 g for 1 h. Membranes were washed twice with lysis buffer before being resuspended in solubilization buffer (50 mM Tris-HCl pH 7.4, 10 % [w/v] glycerol, 250 mM sucrose, 5 mM 6-aminocaproic acid, 5 mM benzamidine, 1 mM PMSF) and solubilized by the addition of glycol-diosgenin (GDN) to 1 % (w/v) and mixing for 1 h at room temperature. Insoluble material was removed by centrifugation at 184,000 g for 45 min and solubilized membranes were loaded onto a 5 ml HisTrap HP column (GE Healthcare) equilibrated with buffer A (solubilization buffer with 20 mM imidazole, 300 mM sodium chloride, and 0.02 % [w/v] GDN). The column was washed with 5 column volumes of buffer A, and ATP synthase was eluted with 3 column volumes of buffer B (buffer A with 200 mM imidazole). Fractions containing ATP synthase were pooled and concentrated prior to being loaded onto a Superose 6 increase 10/300 column (GE Healthcare) equilibrated with gel filtration buffer (20 mM Tris-HCl pH 7.4, 5 mM MgCl_2_, 10 % [w/v] glycerol, 150 mM sodium chloride, 5 mM 6-aminocaproic acid, 5 mM benzamidine, 0.02 % [w/v] GDN). The peak corresponding to *Bacillus* PS3 ATP synthase was pooled and concentrated to ~10 mg/ml prior to storage at -80 °C.

#### Cryo-EM and image analysis

Prior to grid freezing, glycerol was removed from samples with a Zeba spin desalting column (Thermo Fisher Scientific). Purified ATP synthase (2.5 μL) was applied to homemade nanofabricated EM grids ^62^ consisting of a holey layer of gold ^63,64^ that had been glow-discharged in air for 2 min. Grids were then blotted on both sides in a FEI Vitrobot mark III for 26 s at 4 °C and ~100 % RH before freezing in a liquid ethane/propane mixture ^65^. Cryo-EM data were collected with a Titan Krios G3 electron microscope (Thermo Fisher Scientific) operated at 300 kV equipped with a Falcon 3EC direct detector device camera automated with *EPU* software. Data were recorded as 60 s movies at 2 seconds per frame with an exposure rate of 0.8 electron/pixel/second, and a calibrated pixel size of 1.06 Å.

All image processing steps were performed in *cryoSPARC v2* ^66^ unless otherwise stated. 10,940 movies were collected. Movie frames were aligned with an implementation of *alignframes_lmbfgs* within *cryoSPARC v2* ^67^ and CTF parameters were estimated from the average of aligned frames with *CTFFIND4* ^68^. 1,866,804 single particle images were selected from the aligned frames with *Relion 2.1* ^69^ and beam-induced motion of individual particles corrected with an improved implementation of *alignparts_lmbfgs* within *cryoSPARC v2* ^67^. A subset of 1,238,140 particle images were selected by 2D classification in *cryoSPARC v2*. After initial rounds of ab-initio 3D classification and heterogeneous refinement, three classes corresponding to three main rotational states of the enzyme were identified, containing 405,432, 314,448, and 175,694 particles images (Figure 1 - figure supplement 2). These 3D classes were refined with non-uniform refinement to overall resolutions of 2.9 Å, 2.9 Å and 3.1 Å, respectively, with the F_1_ region reaching higher resolution than the F_O_ region of the complex as seen from estimation of local resolution (Figure 1 - figure supplement 3). Masked refinement with signal subtraction (focused refinement) ^33^ around subunits *ab_2_c_10_δ* excluding the detergent micelle improved the map quality of the membrane-embedded region as well as the peripheral stalk for all three classes. The membrane-embedded region (subunits *ac_10_* and transmembrane α-helices of the *b*-subunits) was improved further by focused refinement with particle images from all three classes, yielding a map at 3.3 Å resolution. All Fourier shell correlation (FSC) curves were calculated with independently refined half-maps and resolution was assessed at the 0.143 criterion with correction for the effects of masking maps. For illustration purposes, composite maps for each of the three rotational states were generated by combining the F_1_ region of the maps from non-uniform refinement, the peripheral stalk region from the maps obtained with focused refinement of subunits *ab_2_c_10_δ*, and the map from focused refinement of the membrane-embedded region. Specifically, each map was multiplied by a mask surrounding the region of interest and the resulting maps were adjusted to similar absolute grey scale by multiplying with a constant with *relion_image_handler* before being merged with the maximum function volume operation in *UCSF Chimera* ^70^. These composite maps were not used for model refinement.

#### Model building and refinement

Atomic models for subunits *α_3_β_3_γεδ* from all three rotational states were built with *Coot* ^71^ into the maps of the intact complex from non-uniform refinement using PDB 4XD7 ^19^ and PDB 6FKF ^15^ as initial models for subunits *α_3_β_3_γε* and subunit *δ*, respectively. Subunits *ac_10_* and the membrane-embedded regions of subunits *b_2_* were built *de novo* in the 3.3 Å map of the membrane-embedded region of the complex from focused refinement. Backbone models of the soluble region of subunits *b_2_* for all three conformations were built with the maps from focused refinement of the peripheral stalk. Models were refined into their respective maps with *phenix.real_space_refine* ^72^ using secondary structure and geometric restraints followed by manual adjustments in *Coot* (Table. S1). The quality of the models was evaluated by *MolProbity* ^73^ and *EMRinger* ^74^. To generate full models for all three rotational states, the model of subunits *ac_10_* and the membrane region of subunit *b_2_* were fit into the full maps of each conformation as three rigid bodies (*a*, *c_10_*, and *b_2_* membrane region) with *phenix.real_space_refine*. For class 1 and 3, the backbone models of the soluble region of subunit *b_2_* did not fit the full maps well, and thus the fit was improved by molecular dynamics flexible fitting (MDFF) ^75^. The final composite model for each rotational state was generated by combining the models of subunits *α_3_β_3_γεδ*, the rigid body refined subunits *ac_10_* and membrane region of *b_2_*, and the backbone model of the soluble region of *b_2_*. Figures and movie were generated with Chimera ^70^ and ChimeraX ^43^.

### Figure Captions

**Figure 1 - figure supplement 1.**
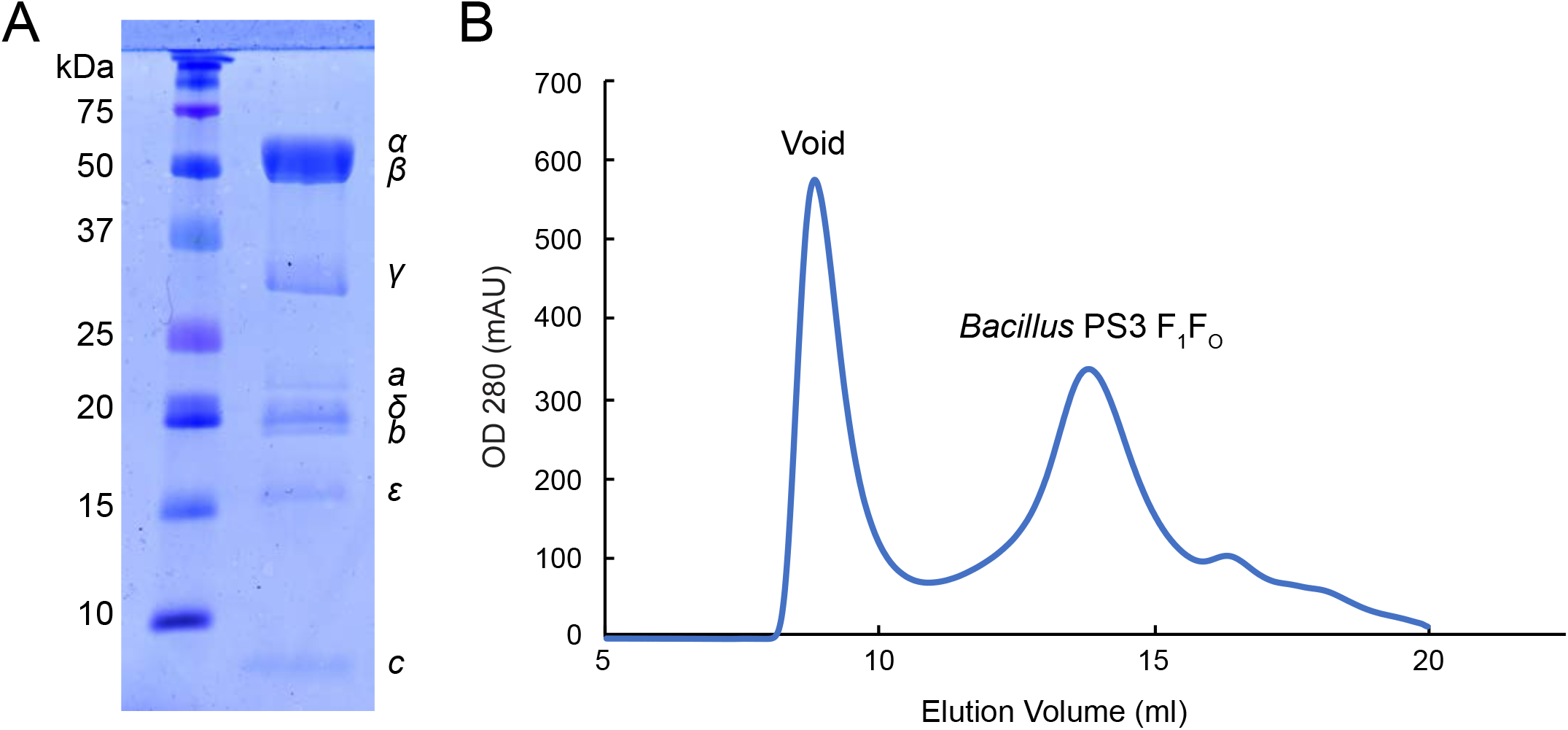
*Bacillus* PS3 ATP synthase purification. **A,** SDS-PAGE of GDN solubilized *Bacillus* PS3 ATP synthase. **B,** Size-exclusion chromatography of *Bacillus* PS3 ATP synthase.

**Figure 1 - figure supplement 2.**
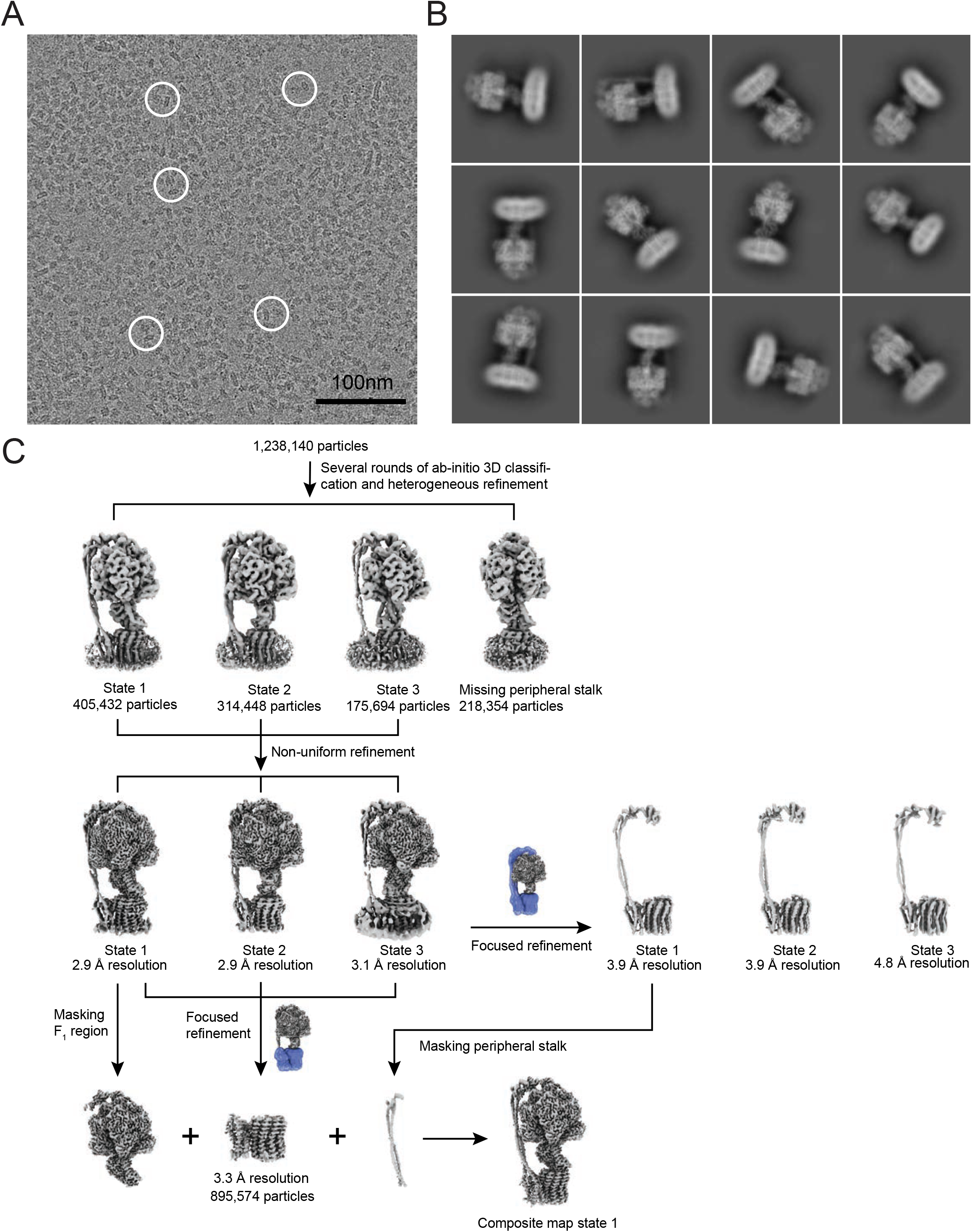
Cryo-EM image processing. **A,** Example micrograph of *Bacillus* PS3 ATP synthase. **B,** Representative 2D class averages of *Bacillus* PS3 ATP synthase. **C,** Image processing workflow.

**Figure 1 - figure supplement 3.**
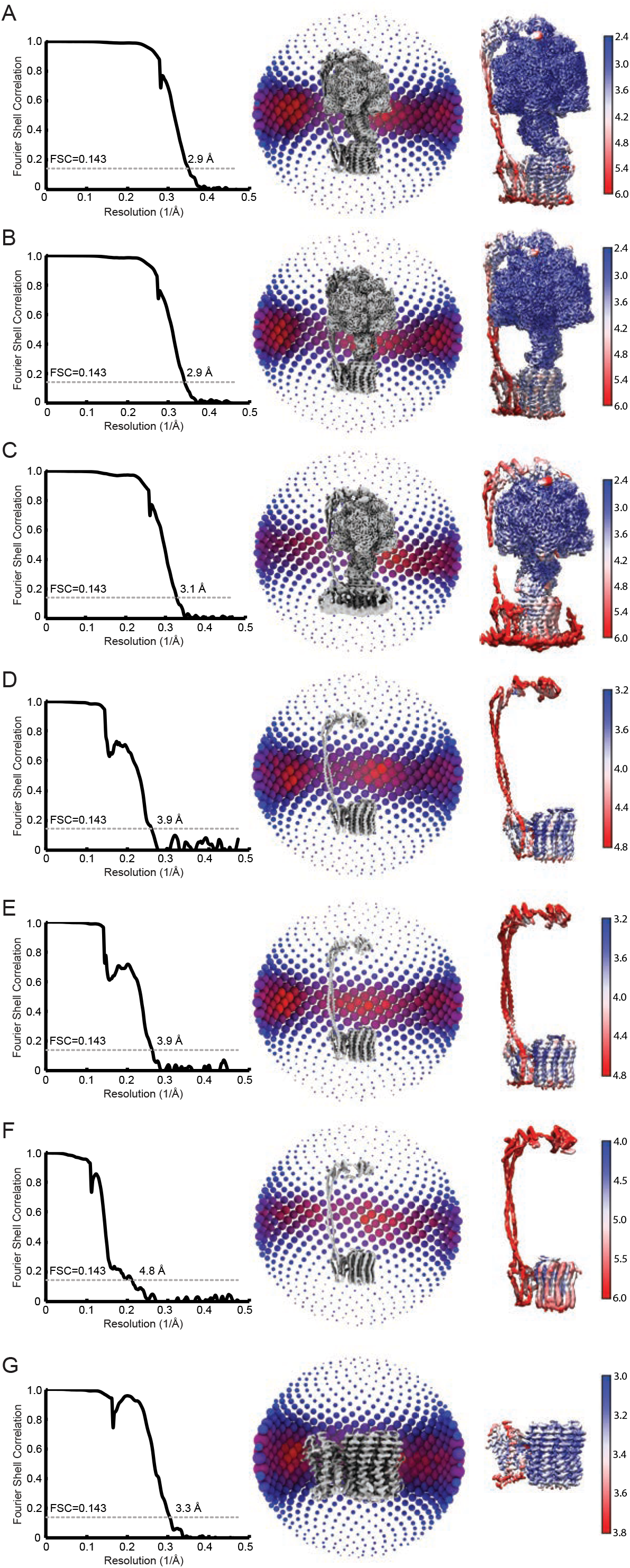
FSC, orientation distribution, and local resolution of the cryo-EM maps used to build atomic models. **A to C,** Full maps of *Bacillus* PS3 ATP synthase class 1, 2, and 3. **D to F,** Focused refinement maps of class 1, 2 and 3 including subunit *δ* and the F_O_ subunits. **G,** Focused refinement map of the membrane bound region only.

**Figure 1 - figure supplement 4.**
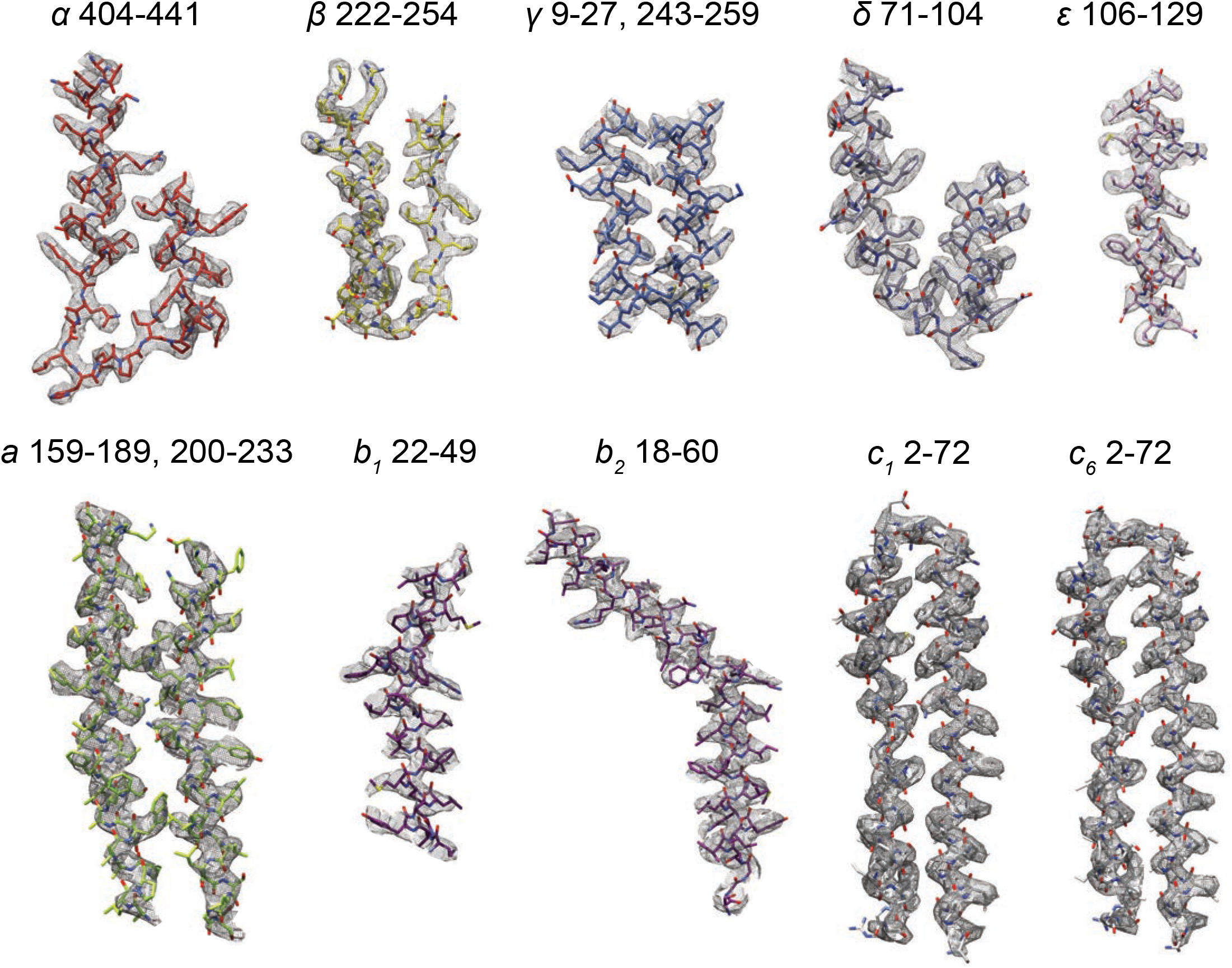
Examples of atomic models from subunits built in the experimental cryo-EM maps.

**Figure 1 - figure supplement 5.**
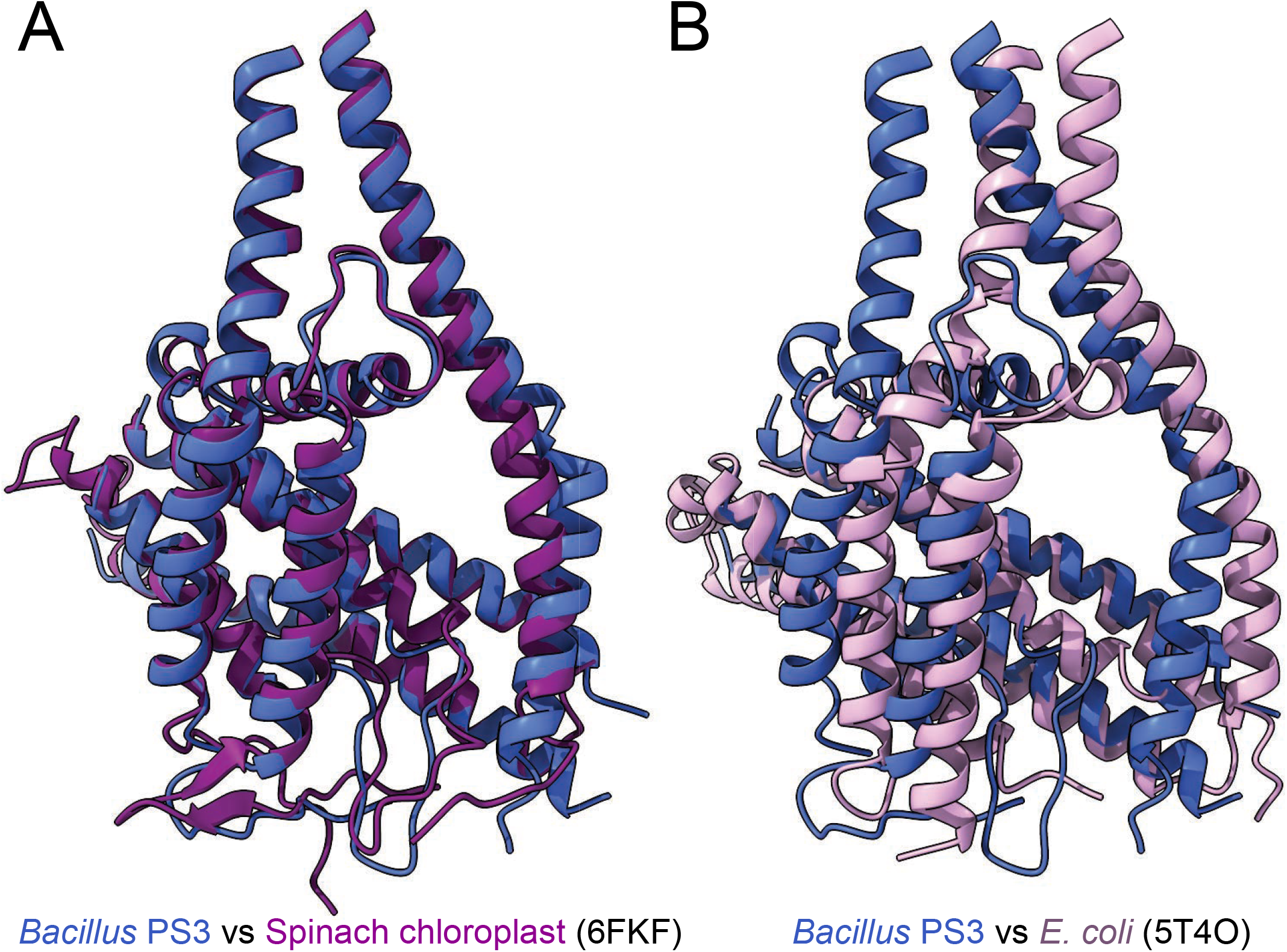
Overlay of *Bacillus PS3* ATP synthase subunits *ab_2_* with spinach chloroplast ATP synthase subunit abb*′* PDB 6FKF ^15^ (**A**) and *E. coli* ATP synthase subunits ab_2_ PDB 5T4O ^25^ (**B**).

**Figure 4 - figure supplement 1.**
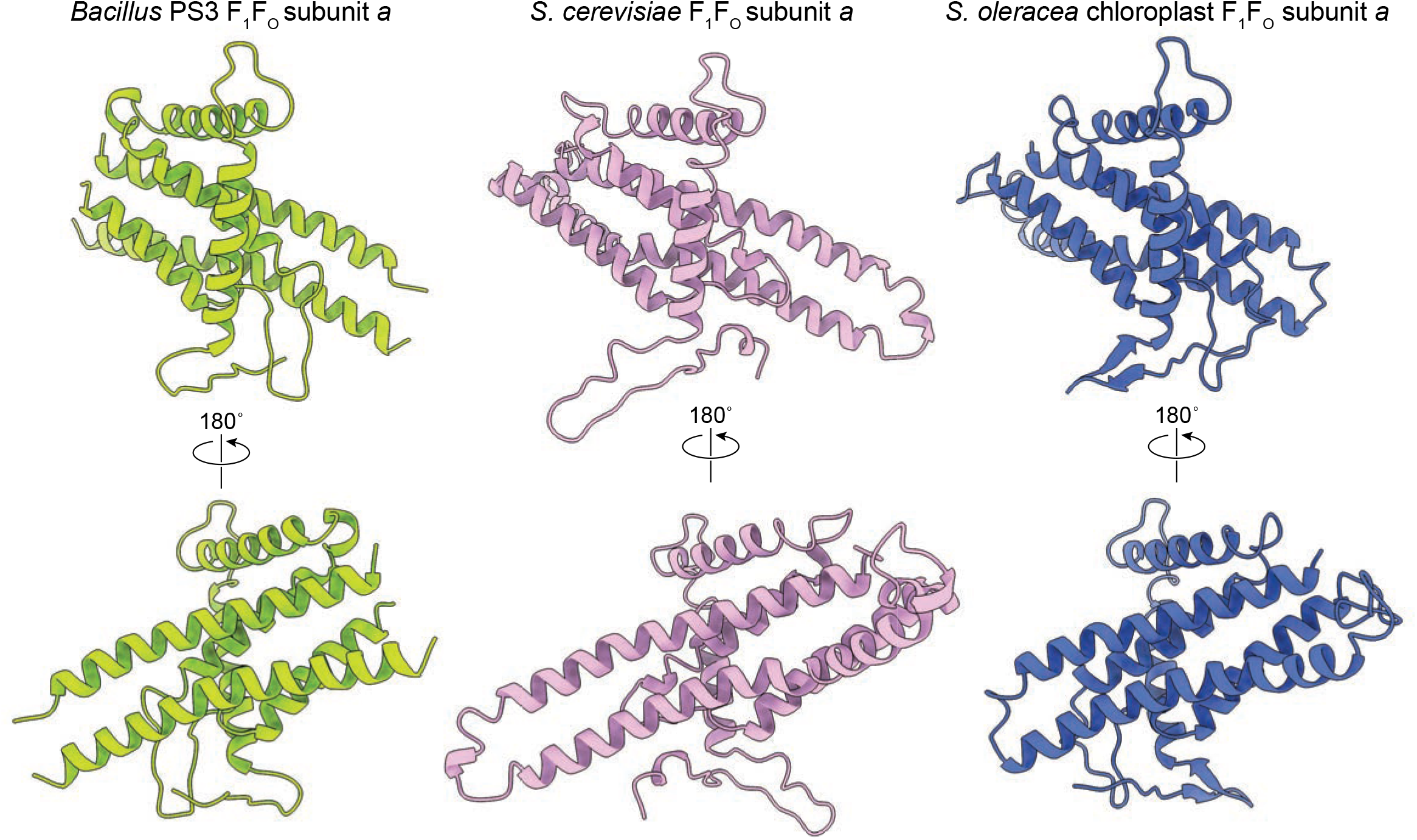
Comparison of subunit *a* structures from different organisms. **A,** *Bacillus* PS3 ATP synthase. **B,** *S. cerevisiae* mitochondrial ATP synthase (PDB 6B2Z ^12^). **C,** *S. oleracea* chloroplast ATP synthase (PDB 6FKF ^15^).

**Figure 4 – figure supplement 2.**
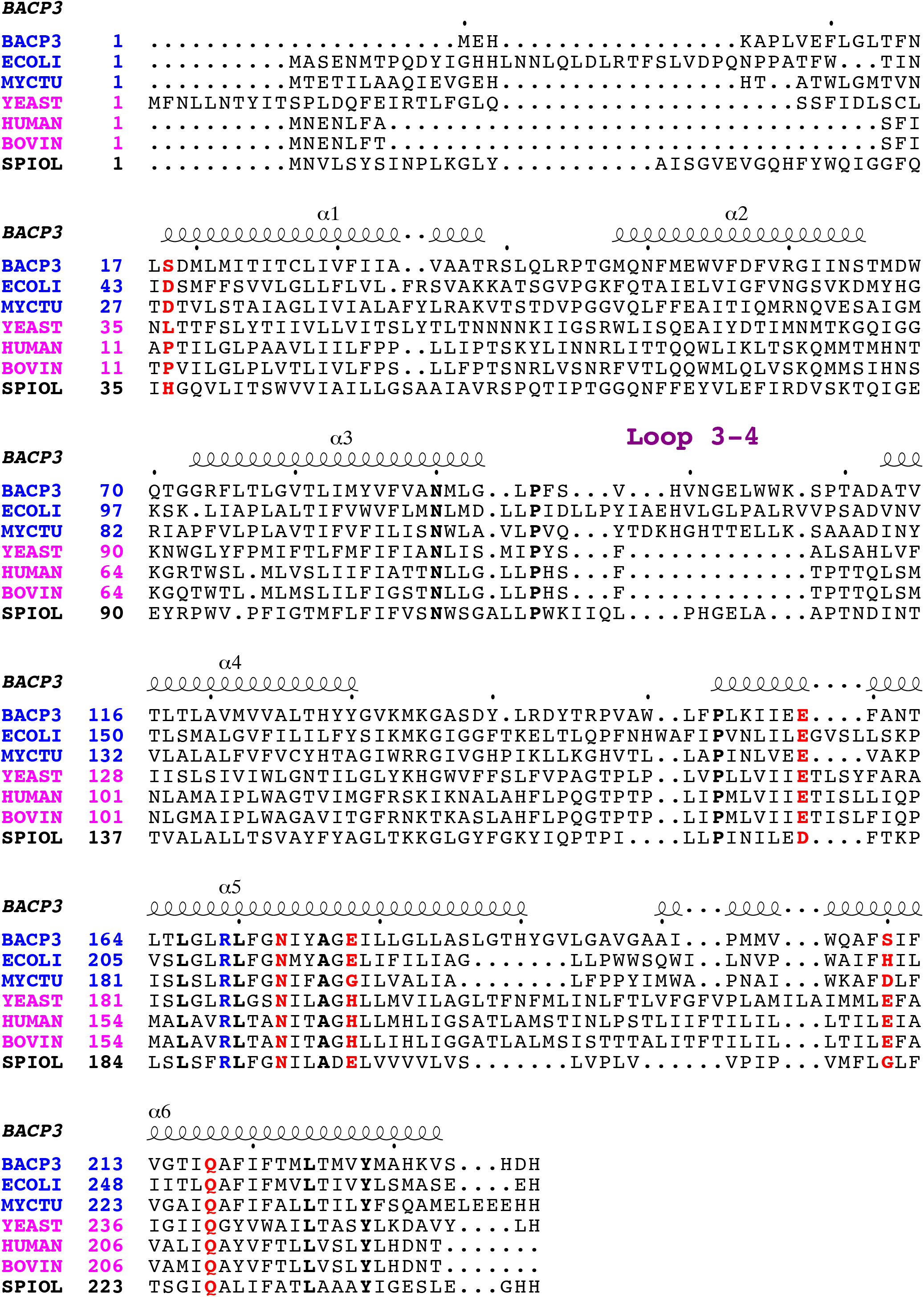
Multiple sequence alignment of subunit *a*. Sequences are from bacteria (blue), mitochondria (pink), and chloroplast (black), including *Bacillus* PS3 (BACP3), *E. coli* (ECOLI), *Mycobacterium tuberculosis* (MYCTU), *S. cerevisiae* (YEAST), *Homo sapiens* (HUMAN), *Bos taurus* (BOVINE), and *Spinacia oleracea* (SPIOL). The functionally important Arg residues for proton translocation are in blue. Other important residues for proton translocation identified by mutagenesis in *E. coli* ATP synthase are in red. Strictly conserved residues are in bold.

**Figure 4 - figure supplement 3.**
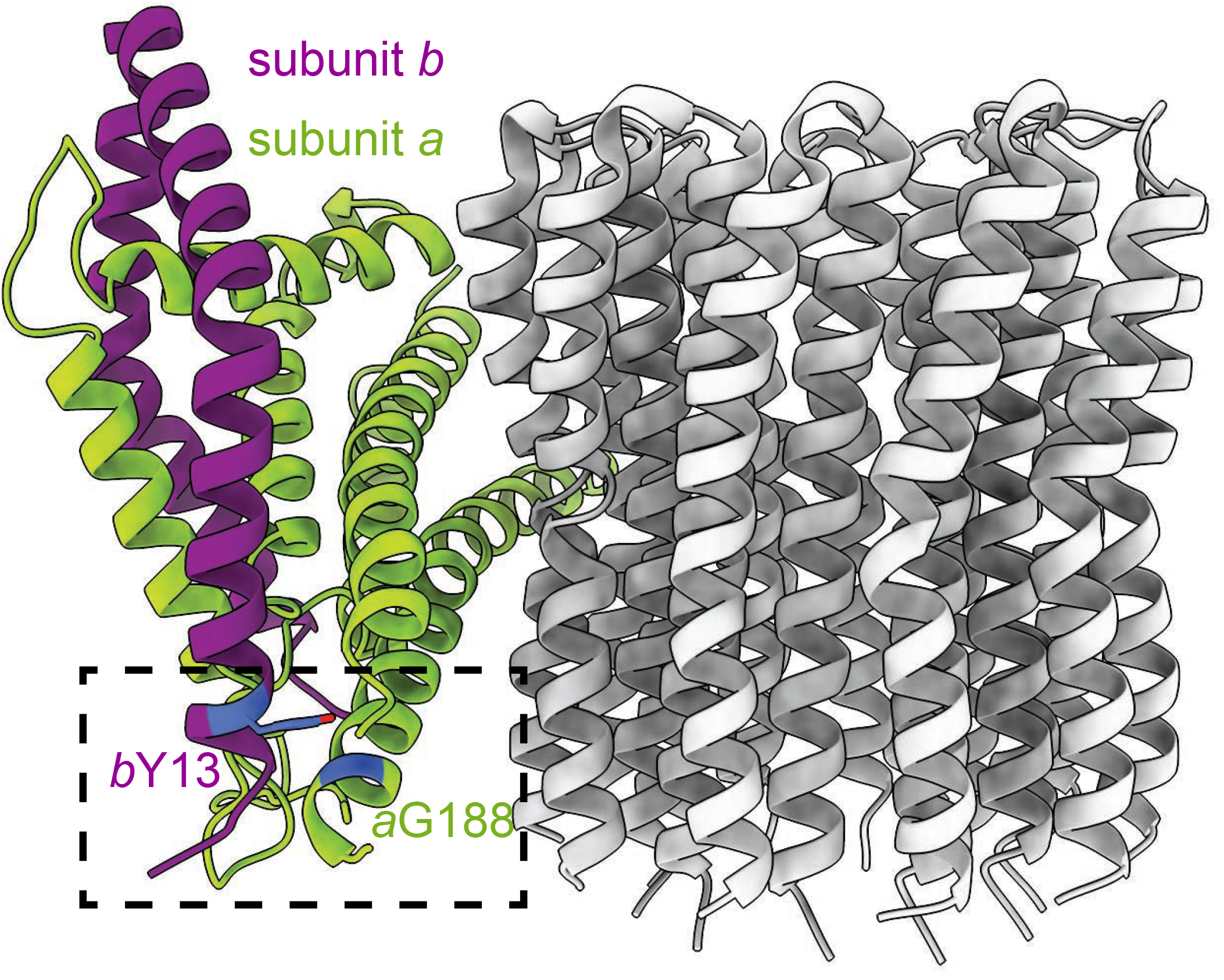
Positions of subunit *b* Tyr 13 and subunit *a* Gly 188. Side view of the F_O_ subunits. The residues of interest are inside the dashed box and are coloured in blue.

**Table 1.** Cryo-EM data acquisition, processing, atomic model statistics and map/model depositions a. Cryo-EM data acquisition and image processing. b. Map and model statistics. c. Deposited maps and associated coordinate files

| **Data Collection** |  |
| --- | --- |
| Electron Microscope | Titan Krios |
| Camera | Falcon 3EC |
| Voltage (kV) | 300 |
| Nominal Magnification | 75,000 |
| Calibrated physical pixel size (Å) | 1.06 |
| Total exposure (e/Å<sup>2</sup>) | 42.7 |
| Exposure rate (e/pixel/s) | 0.8 |
| Number of frames | 30 |
| Defocus range (µm) | 0.5 to 3.8 |
| **Image Processing** |  |
| Motion correction software | *cryoSPARC v2* |
| CTF estimation software | *CTFFIND 4* |
| Particle selection software | *Relion 2.1* |
| Micrographs used | 10,940 |
| Particle images selected | 1,866,804 |
| 3D map classification and refinement software | *cryoSPARC v2, Relion 2.1* |

**b. Map and model statistics.**
| EM maps | Intact class 1 | Intact class 2 | Intact class 3 | Focused Fo/stalk class 1 | Focused Fo/stalk class 2 | Focused Fo/stalk class 3 | Focused Fo |
| --- | --- | --- | --- | --- | --- | --- | --- |
| Particle images contributing to maps | 405,432 | 314,448 | 175,694 | 405,432 | 314,448 | 175,694 | 895,574 |
| Applied symmetry | C1 | C1 | C1 | C1 | C1 | C1 | C1 |
| Applied B-factor (Å²) | -124.7 | -122.2 | -111.9 | -169.2 | -153.1 | -175.7 | -155.2 |
| Global resolution (FSC = 0.143, Å) | 2.9 | 2.9 | 3.1 | 3.9 | 3.9 | 4.8 | 3.3 |
| Model Building | F<sub>1</sub> class 1 |  | F<sub>1</sub> class 2 |  | F<sub>1</sub> class 3 |  | F<sub>0</sub> |
| Modeling software | Coot, Phenix |  |  |  |  |  |  |
| Residue numbers | 3501 |  | 3497 |  | 3499 |  | 993 |
| RMS bond length (Å) | 0.0066 |  | 0.0079 |  | 0.0093 |  | 0.0084 |
| RMS bond angle (°) | 1.28 |  | 1.32 |  | 1.38 |  | 1.48 |
| Ramachandaran outliers (%) | 0.17 |  | 0.12 |  | 0.14 |  | 0.00 |
| Ramachandran favoured (%) | 96.03 |  | 95.73 |  | 95.13 |  | 97.40 |
| Clashscore | 5.69 |  | 5.97 |  | 5.95 |  | 14.25 |
| MolProbity score | 1.58 |  | 1.62 |  | 1.66 |  | 1.78 |
| EMRinger score | 4.50 |  | 4.44 |  | 4.49 |  | 1.68 |
| Ligand | 1 Pi<br>1 Mg-ADP<br>3 Mg-ATP |  | 1 Pi<br>1 Mg-ADP<br>3 Mg-ATP |  | 1 Pi<br>1 Mg-ADP<br>3 Mg-ATP |  |  |

**c. Deposited maps and associated coordinate files**
| **Maps** | **EMDB code** | **Associated PDB ID** |
| --- | --- | --- |
| Intact class 1 | EMD- |  |
| Intact class 2 | EMD- |  |
| Intact class 3 | EMD- |  |
| Focused Fo/stalk class 1 | EMD- |  |
| Focused Fo/stalk class 2 | EMD- |  |
| Focused Fo/stalk class 3 | EMD- |  |
| Focused Fo | EMD- |  |

